# The Rapid-Tome, a 3D-Printed Microtome, and an Updated Hand-Sectioning Method for High-Quality Plant Sectioning

**DOI:** 10.1101/2022.10.28.513790

**Authors:** David J. Thomas, Jordan Rainbow, Laura E. Bartley

## Abstract

**Background:** Microscopic analysis of plant anatomy is a common procedure in biology to study structure and function that requires high-quality sections for accurate measurements. Hand sectioning of specimens is typically limited to moderately soft tissue while harder samples prohibit sectioning by hand and/or result in inconsistent thicknesses.

**Results:** Here we present both a clearly described hand-sectioning method and a novel microtome design that together provide the means to section a variety of plant sample types. The described hand-sectioning method for herbaceous stems works well for softer subjects but is less suitable for samples with secondary growth (e.g., wood production). Instead, the “Rapid-Tome” is a novel tool for sectioning both soft and tougher high-aspect-ratio samples, such as stems and roots, with excellent sample control. The Rapid-Tome can be 3D-printed in approximately 13 hours on a mid-quality printer common at university maker spaces. After printing and trimming, Rapid-Tome assembly takes a few minutes with five metal parts common at hardware stores. Users sectioned a variety of plant samples including the hollow internodes of switchgrass (*Panicum virgatum*), fibrous switchgrass roots containing aerenchyma, and woody branches of eastern red cedar (*Juniper virginiana*) and American sycamore (*Platanus occidentalis*). A comparative analyses with Rapid-Tome-produced sections readily revealed a significant difference in seasonal growth of sycamore xylem vessel area in spring (49%) vs. summer (23%). Additionally, high school students with no prior experience produced sections with the Rapid-Tome adequate for comparative analyses of various plant samples in less than an hour.

**Conclusions:** The described hand-sectioning method is suitable for softer tissues, including hollow-stemmed grasses and similar samples. In addition, the Rapid-Tome provides capacity to safely produce high-quality sections of tougher plant materials at a fraction of the cost of traditional microtomes combined with excellent sample control. The Rapid-Tome features rapid sectioning, sample advancement, blade changes, and sample changes; is highly portable and can be used easily with minimal training making production of thin sections accessible for classroom and outreach use, in addition to research.

## Background

Plant anatomy is a wondrous micro-verse to explore, rich with information on cellular structure and function. Plant anatomy is not only beautiful to observe, but through combining anatomical phenotyping with genetic analyses, the ability to optimize plant anatomy for various desired functionality draws closer. Great efforts have been devoted to understanding the impact of cell division, expansion, and identity on shape formation of plant organs [1]. Increasing our understanding of the connection between molecular mechanisms with plant form and function may lead to synthetic morphogenesis [2]. The availability of tools that facilitate generating anatomical phenotype data is crucial.

Investigations of genetic regulation of morphogenesis require characterization of phenotypes, including in forward genetic screens [3]. For example, over 20 years ago, the field of secondary cell wall synthesis was propelled by the identification of irregular xylem mutants through screening an Arabidopsis population via microscopic examination of stem sections [4]. More recently, a forward genetic screen of abnormal vasculature in cross sections of *Brachypodium distachyon* identified a mutant phenotype caused by a premature stop codon in *BdERECTA* that impacts regulation of vascular bundle development and may impact function [5]. In another recent example, laser ablation tomography (LAT) was applied to screen for root anatomy phenotypes in cross sections of a genome wide association panel of maize (*Zea mays)* to identify multiseriate cortical sclerenchyma correlated with greater root tip lignin concentrations and bending strength [6]. Emerging image analysis methods that utilize machine learning approaches will further accelerate studies and modeling of anatomy [7]. These examples all rely on high-quality sections to make accurate measurements and identify phenotypes.

Individual cell traits visible in sectioned plant tissues, i.e. dimensions or cell wall thickness, can provide insight into the functional performance of different cell types. To accurately measure these traits, sections that are flat, consistent thickness, and thin enough to place on a microscope slide are required. Section integrity is the most impactful variable towards obtaining high-quality images, and therefore, high-accuracy measurements at the image analysis stage [8]. Section thickness is a major constraint for transmitted light microscopy. The requirements for high-quality sections can depend on the nature of the subject material. The hardness and thickness of the material determine the best sectioning approach and the difficulty of obtaining thin, high-quality sections. Rigid, sclerenchymatic cells of vascular bundles and interfascicular fibers are often juxtaposed with softer cells [9]. Cell type heterogeneity increases the difficulty to obtain high-quality sections by hand. Furthermore, the presence of weaker cells intermixed with tough cells, or a hollow specimen, is also problematic because the risk of crushing the sample when clamping into place in a microtome unless strengthened by embedding the plant material in a hard material such as wax or resin.

Options for producing plant sections for light microscopy range in technical difficulty, sample preparation, learning required, and cost (Table 1). Hand sectioning with only a razor blade requires the least investment and is valuable for early investigations and mostly limited to softer tissues. However, section thickness is difficult to control and can vary among researchers. Surprisingly, we were unable to locate literature on the details of a hand sectioning method. Hand sectioning can be used on soft tissues, like young roots or leaves. If more control is needed, such soft tissues can also be gently embedded in low-melt temperature agarose for relatively easy sectioning with a vibratome [10]. Hand sectioning and vibratome sectioning can both be completed on the same day as sampling. However, these methods can be difficult to apply to tough stems, which require great control over sample and blade advancement to section thinly.

**Table 1.** Approaches and tools to prepare plants for imaging in cross section.

| Method | • Section quality • Accuracy<sup>a</sup> | Sample preparation | Required materials | Learning curve | Benchtop required | Portable | Cost |
| --- | --- | --- | --- | --- | --- | --- | --- |
| **Hand sectioning** |  |  |  |  |  |  |  |
| Chopping on bench | • Poor<br>• Poor | None | Razor blades | Low | Yes | No | \$ |
| Method described herein | • High<br>• High | None | • Razor blades<br>• Parafilm | Medium - high | No | Semi | \$ |
| **Microtomes** |  |  |  |  |  |  |  |
| Hand microtomes | • High<br>• High | Cut to fit clamps | Razor blades | Medium | Depends on model | Semi | \$\$ |
| Benchtop microtomes | • High<br>• High | Days to for dehydration and embedding | • Embedding materials and equipment<br>• May require electricity | Medium - high | Yes | No | \$\$\$ |
| Vibratome | • High<br>• Medium | Hours to days for embedding | • Embedding materials and equipment<br>• Electrical outlet | Medium - high | Yes | No | \$\$\$ |
| Rapid-Tome | • High<br>• High | None | • Access to 3D-printer<br>• Razor blades<br>• Nuts and bolts<br>• Teflon dry lube<br>• Adhesive foam | Low | No | Field capable | \$ |
| Laser Ablation Tomography<sup>c</sup> | • High<sup>b</sup><br>• High<sup>b</sup> | None | • Laser<br>• Microscope<br>• Camera | High | Yes | No | \$\$\$\$ |

a- Accuracy is defined as the ability to target and produce sections of a specific region if interest
b- Material is blasted away by the laser and exposed surface imaged to generate digital sections as the sample is destroyed.
c- Schneider, H. M., Strock, C. F., Hanlon, M. T., Vanhees, D. J., Perkins, A. C., Ajmera, I. B., Sidhu, J. S., Mooney, S. J., Brown, K. M., & Lynch, J. P. (2021). Multiseriate cortical sclerenchyma enhance root penetration in compacted soils. *Proceedings of the National Academy of Sciences*, 118(6).

Benchtop microtomes which are often used for obtaining very thin, uniform sections typically require sample dehydration and embedding, which can be time-consuming depending on the sample type and method [11]. Though time consuming, resin embedded samples can be tightly clamped into place for obtaining very thin sections (∼5-50 μm) with a microtome or ultra-microtome. Paraffin wax embedding, commonly used for plant material [12], requires handling of molten wax and hours to days for wax infiltration of the sample. Another typical embedding medium, HistoResin, is limited to rather small blocks, preventing the analysis of larger plant subjects. Furthermore, it involves a medium to high learning curve and a benchtop microtome [13-15]. A major consideration with this process is that embedding in resin or wax requires the samples to be dehydrated first, which in turn greatly influences the success of resin infiltration and the quality of the sections obtained. Thick samples and those with highly dense and numerous cells take longer to both completely dehydrate and resin infiltrate. Together, the time necessary for dehydration and resin infiltration may at times be on the order of weeks. In addition, pilot investigations and multiple trials are typical to determine the required length of dehydration and time to achieve complete resin infiltration which vary with the type of resin and anatomical features such as high starch content or thick cell walls [16, 17].

The progression of additive manufacturing technology and the decreasing cost of three dimensional (3D) printers have created an opportunity for new tools that can be printed on site and utilized in research labs and classrooms alike. A repository for downloadable 3D models for assisting in plant research was recently made available as a community resource in 2020 at “3DPlantPhenomics.org” [18]. Several recent studies utilized 3D-printing to enhance plant tissue sectioning methods or throughput [19]. For soft tissues, Atkinson et al. created a 3D-printed embedding mold to embed plant roots in agarose for sectioning. Another group created a 3D-printed sectioning tool produce sections of plant material, though this tool has no sample advancement mechanism to control section thickness [20].

Here, with the goal of facilitating inexpensive and rapid analysis of plant organ cellular morphology, we present two sectioning methods for elongated, harder plant materials, like crop roots and stems/culms. The second of these, a 3D-printable microtome, the Rapid-Tome, takes advantage of additive manufacturing to generate a device that can produce high-quality sections with a low learning curve suitable for use in the field and with novices.

## Results and Discussion

The most common approach to sectioning fresh tissue in classrooms and the laboratory setting is hand sectioning. Here we present a detailed hand sectioned method that is highly effective with fresh and soft plant tissues such as herbaceous stems, petioles, roots, etc. It is can, however, be variable among users and difficult to generate complete sections. Further, when sectioning harder samples that are highly lignified or woody, the hand-sectioning method is not easily applied, and the use of pressure required to cut with a razor blade by hand can be dangerous. To address these concerns, we present a novel, 3D-printed, hand-held microtome that can produce high-quality sections even in the hands of a novice.

### Hand sectioning

The hand sectioning method is described in detail in Figure 1, including an approach to section hollow stems that collapse (Figure 1K). Figure 2A shows the quality of section that this method can produce. Both images are less than a single cell thick (< 100 μm) and exceed the quality required to accurately measure anatomical traits such as cell wall thickness and lumen diameter or area. When the section is less than 1 cell thick, there are no overlapping cell walls from overlying or underlying cells to convolute the field of view so the cell’s walls can be identified with confidence. The limitation here is with the microscope and magnification rather than the section. However, tougher subjects that in the case of the internodes contain more lignin in the sclerenchyma fibers prove difficult to hand section and often break apart (Figure 2B and 2C). Furthermore, the tough cell layers (e.g., in the sclerenchyma) are often marked with chatter marks from the blade making inconsistent contact (Figure 2C, black arrowheads), and in some cases will cause the tissue to break (Figure 2C, white arrowhead). In such situations, and to attain more consistent sections, we recommend the use of a sectioning tool.

**Figure 1.**
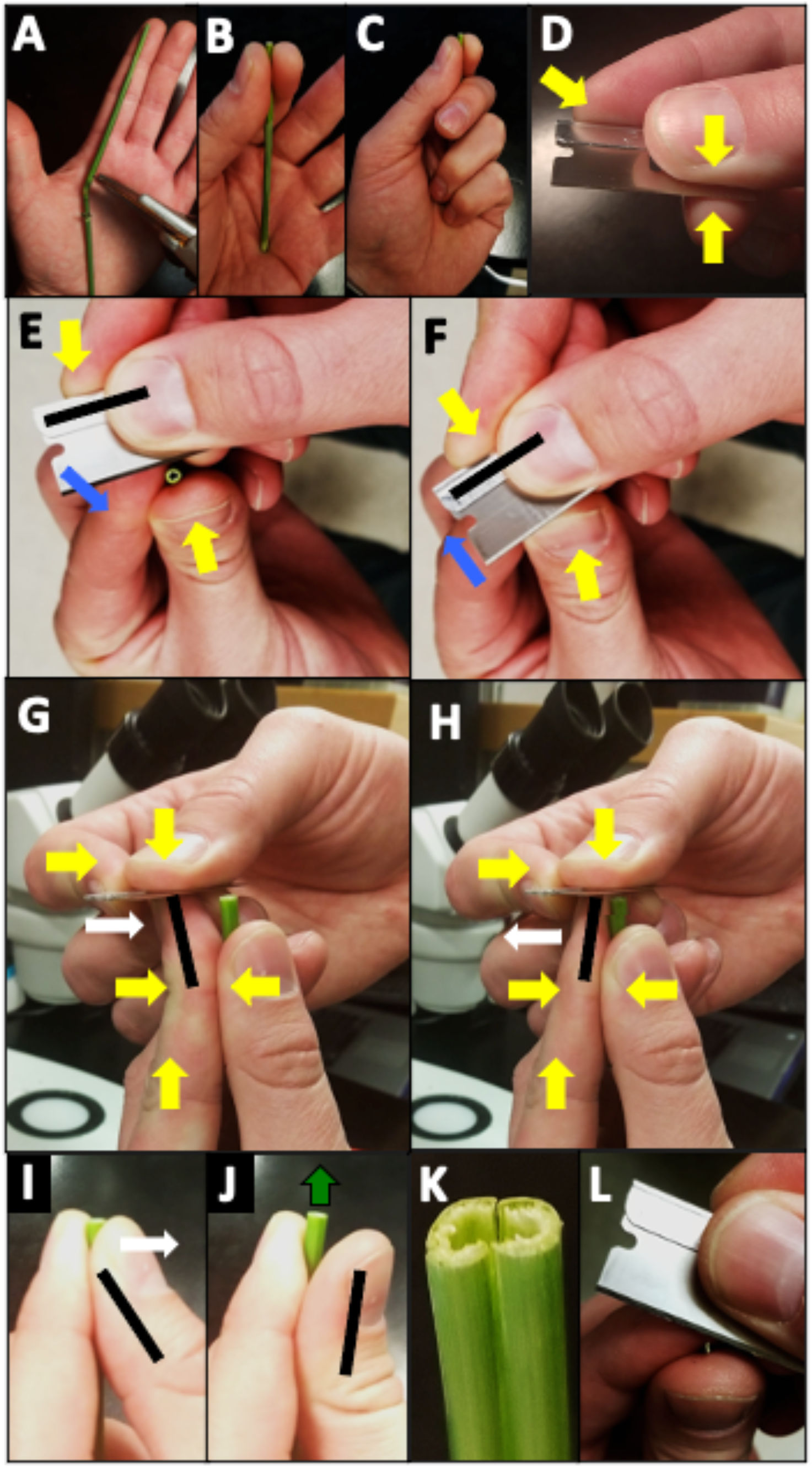
Hand sectioning of a grass internode with a single-edge razor blade. (A) Cut the stem segment to length to match the size of the sectioner’s hand as shown. Stem segment is shown being sectioned by a right-handed person. (B) The stem is held between the thumb and forefinger and pressed into the palm gently to further stabilize in the left hand. (C) Fingers are wrapped around the stem. (D) Yellow arrows represent static pressure applied by fingers to hold the blade firmly in place. (E and F) The blade is held in the dominant hand while the plant stem is in the nondominant hand. The blade is moved forward by the movement of the dominant-hand’s forefinger tip, which is slightly bent to rotate the blade slightly as the wrist flexes (Blue arrows). (G and H) The flat surface of the blade in the dominant hand is gently pressed downward against the forefinger of the nondominant hand. The blade advances to take sections and then retreats with the movement of the nondominant-hand forefinger and dominant hand wrist. Once started, this motion should be repeated without separating the hands for several sections. (E and F) Blue arrow represents blade movement. Black bars (E-J) represent the orientation of the finger or blade before and after pivoting. (G-I) White arrows represent finger movements. (J) Green arrow represents stem movement upwards by the subtle flexing of the nondominant thumb backwards. Alternatively, the thumb and forefinger can gently slide downward towards the palm to expose more stem for sectioning. (K) A crushed or collapsed stem can be separated to isolate a single shard and section starting with the short edge shown in (L). This is a very slow and gentle movement to prevent cutting a finger.

**Figure 2.**
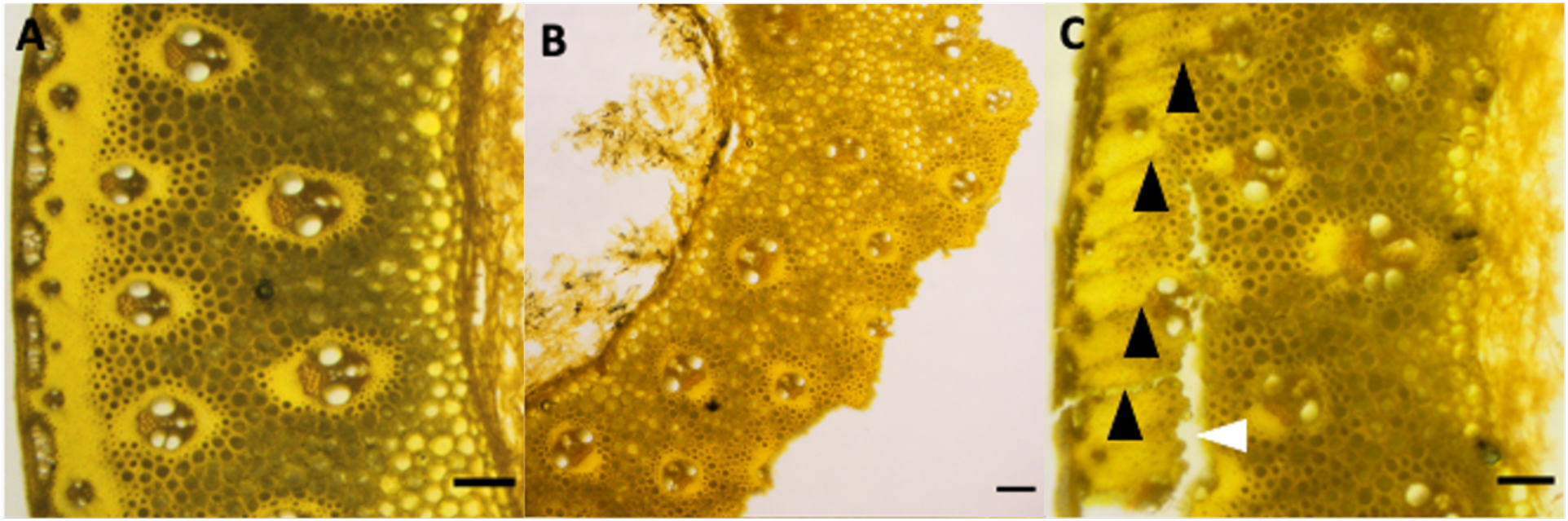
The hardness and heterogeneity of the plant material reduces the quality of sections easily obtained with the hand sectioning method. Samples are switchgrass internode cross sections stained with acriflavine hydrochloride. (A) Adequate quality for image analysis. (B) Tougher samples are not able to be sectioned well with the hand sectioning method and often produce cut artifacts where the sample is pulled apart as the tough sclerenchyma contacts the blade, leading to incomplete sections with entire tissues absent.(C) Blade chatter marked with black arrowheads can interfere with image analysis due to uneven section thickness that causes uneven focal planes, and in some cases causes a full break in the sample (white arrowhead). All scale bars are 100 microns.

### The Rapid-Tome

The Rapid-Tome is a tool for producing high-quality sections from high-aspect ratio plant subjects, such as stems, petioles, and roots in a quick and safe way that can be readily used by a novice. After several rounds of prototypes and design iterations, the final version is shown in Fig. 3 and Additional File 1. The Rapid-Tome is a combination of the gentle and firm support of holding a plant sample in your hand and the highly controlled blade advancement possible with a microtome.

**Figure 3.**
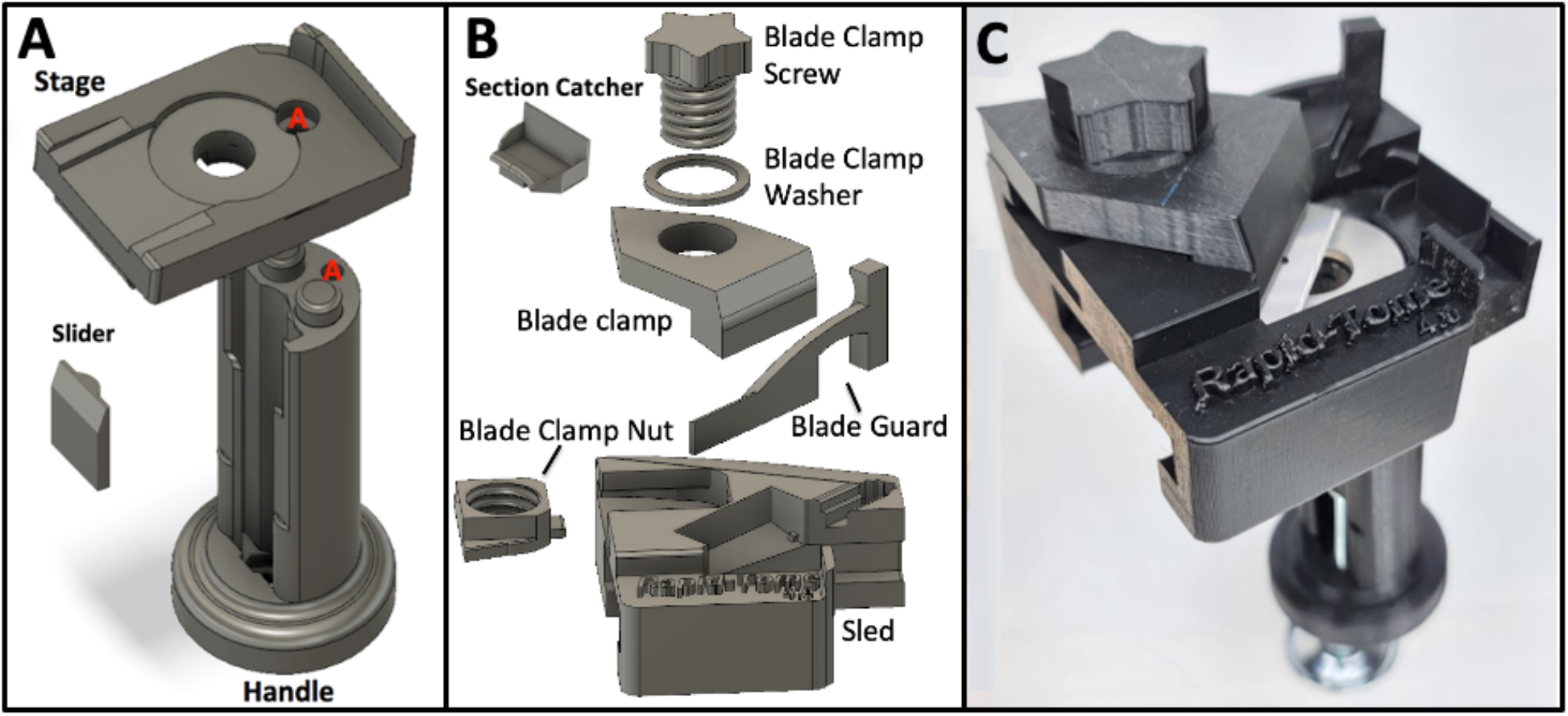
3D-models and fully assembled Rapid-Tome. (A) the handle and stage are connected with a single metal bolt through holes “A” after sliding the slider piece into the vertical grooves form the top. (B) the assembly of the sled includes the attachment of the blade clamp with the blade clamp nut, blade clamp washer, and blade clamp screw. The blade clamp nut should be slid into place first. The blade clamp screw is fed through the top of the blade clamp with the blade clamp washer between. The blade guard should be kept in place over the blade at all times when the sled is removed from the stage. The blade guard is removed during sectioning and stored in the holster contained within the sled. (C) the fully assembled Rapid-Tome with advancement bolt, stage washer, and blade installed.

From our prototyping, we found that the critical points of the Rapid-Tome’s design are the blade clamp and the angle of the stage that holds the blade in a flexed and rigid position. As the sled advances, the peripheral regions of the blade encounters the metal washer. The stage edges that guide the sled advancement is angled against the metal washer which causes the blade to flex upward slightly. The washer surface is at a right angle to the handle. Because of the flexed position of the blade, the cutting edge of the blade is held flat and produces sections with consistent thickness. The clamp holds the blade in a rigid and flexed fashion so that the blade does not travel upward in the z-axis, which produces sections with inconsistent thickness that are wavy or wedged-shaped. Such irregular sections result in inconsistent contact with a coverslip and an inconsistent focal plane and viewing of a large, in-focus field at low magnification. Indeed, when hand sectioning tougher samples such as large and fibrous switchgrass internodes or woody branches of eastern red cedar, control of the blade is difficult to maintain which results in poor section quality and inconsistent section thickness. Because the Rapid-Tome blade is clamped, flexed, and held in place by the sled, the researcher has more control over blade advancement and can proceed slowly and safely. Additionally, the sled and blade clamp keep the operator’s fingers away from the sharp razor and prevent the possibility of injury even if the sample slips and the blade quickly advances.

Another important feature incorporated into the design of the Rapid-Tome is that the sample is held in place against the handle by the researcher’s thumb. Typical microtomes have a hard metal or plastic clamp that tightens with a bolt, requiring the sample to be hard enough and with adequate structural integrity so that it can withstand the force of being clamped into place. The padded skin of the thumb provides firm but delicate pressure with real time modification if necessary that is unrivaled by a clamp. The sample is held steady against the back wall of the handle as the blade cuts in the same direction so that the forces on the specimen are not in opposition. Additionally, since there is no clamp screw or apparatus to remove, the subject can be quickly exchanged to increase throughput.

To demonstrate the utility of the Rapid-Tome, a variety of sample types were sectioned both by experienced researchers and novices. Samples that we tested with the Rapid-Tome include switchgrass internodes (Fig. 4A-C), switchgrass roots (Fig. 4D-F), and small branches from Eastern Red Cedar (*Juniperus virginiana)* (Fig. 4G-I) and American Sycamore (*Platanus occidentalis)* (Fig. 4J-L). Fresh samples can be sectioned very thin even with abundant fibers (f) and with intact subcellular components that include chloroplasts visible within the chlorenchyma (Fig. 4 A-C). Full transverse sections of switchgrass roots stained with toluidine blue contains numerous xylem vessels (x) in the stele and aerenchyma (a) within the cortical parenchyma radiating outward between the pith and the lignified endodermis (e) (Fig. 4D). Higher magnifications bring clearly into view the aerenchyma, outer layers of the root, and a pair of large vessels within the pith with different cell wall thicknesses associated with different cells (Fig. 4E and F).

**Figure 4.**
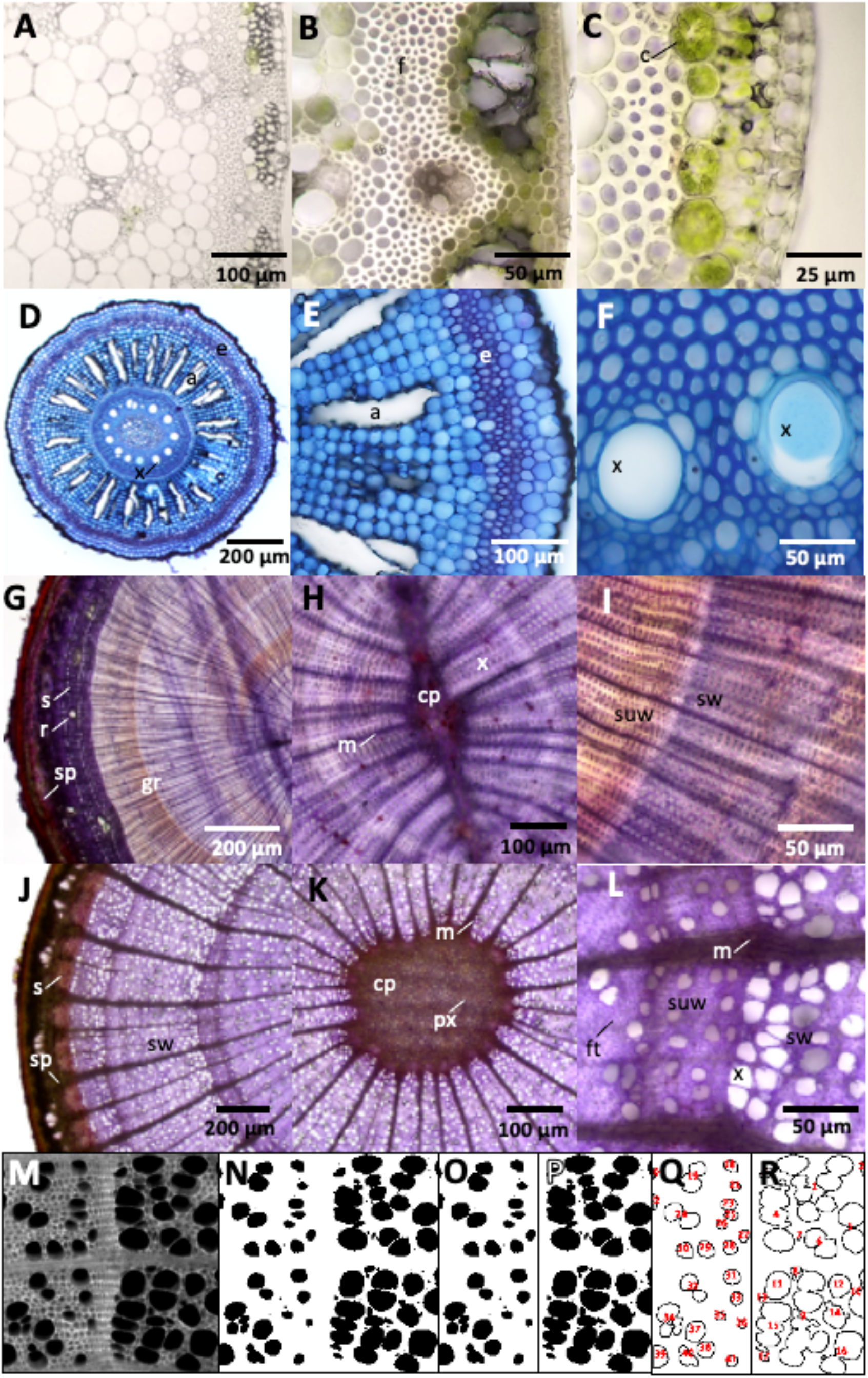
The Rapid-Tome can easily section a variety of plant samples. (A-C) Unstained switchgrass internode. (A) Fresh samples can be sectioned very thin (B) even with abundant fibers and (C) and chloroplasts. (D-F) Switchgrass root stained with toluidine blue. (D) Full transverse section of root with multiple cell types visible. (E) Higher magnification of the root. (F) A pair of large vessels within the pith. (G-L) A series of images for transverse comparative anatomy of conifer and angiosperm twigs. (G-I) Crystal violet-stained eastern red cedar (ERC). (G) Multiple spring and summer growth boundaries are visible. (H) The central pith (cp), surrounding xylem, and radial medullary rays (m) are visible within the first years of growth. (I) Summer wood and spring wood denotes the annual ring. (J-L) Crystal violet-stained American sycamore. (J) Three distinct layers of spring wood are visible. (K) The central pith of the sycamore twig is larger in comparison to ERC. (L) Higher magnification of the annual ring boundary and clearly visible is the large difference in xylem vessel diameter between ERC and Sycamore in (I) and (L). (M-R). Analysis of spring vs. summer wood vessel fraction by area with the high-contrast images of cell wall autofluorescence visible with the GFP filter set. (M) Autofluorescent image. (N) Binary image. (O) Summer wood. (P) Spring wood. (Q and R) Vessel area identified and measured with “analyze particles” tool in FIJI shows that summer wood is 23% vessels by area and 49% in spring wood. All samples sectioned fresh the day of collection. Fibers (f), chloroplasts (c), xylem vessels (x), aerenchyma (a), endodermis (e), growth rings (gr) resin ducts (r), sclerenchyma (s), secondary phloem (sp), summer wood (suw), spring wood (sw), primary xylem (px), fiber tracheid (ft). Scale bars as labelled.

A series of images for transverse comparative anatomy of conifer and angiosperm twigs sectioned to adequate thickness for transmitted light illumination (Fig. 4G-L). Crystal violet-stained eastern red cedar show multiple growth rings (gr) indicated by spring and late growth boundary, resin ducts (r) within the sclerenchyma (s), and secondary phloem (sp) underlying the bark (Fig. 4G). (H) The central pith (cp), surrounding xylem, and radial medullary rays (m) are visible within the first years of growth of the eastern red cedar (Figure 4 H and I). The transition of thinner walled xylem and larger lumen of the xylem in spring wood (sw) compared to thicker walls and smaller lumen in the summer wood (suw) denotes the annual ring (Fig. 4I). (J) Three distinct layers of spring wood are visible crystal violet-stained American sycamore with groups of thick-walled sclerenchyma visible underlying the bark, separated by secondary phloem (Fig. 4J). The central pith of the sycamore twig is larger in comparison to eastern red cedar and is populated with primary xylem (px), surrounded by diffuse porous xylem dissected by radial spaced medullary rays Fig. 4K). Higher magnification of the annual ring boundary reveals summer wood indicated by smaller area ratio of xylem vessels to fiber tracheid (ft) followed by larger ratio in spring wood with visibly large xylem vessels Figure 4 L). Also clearly visible is the large difference in xylem vessel diameter between eastern red cedar and Sycamore in (I) and (L).

In addition, an analysis investigating the difference between xylem vessel area in sycamore (*Platanus occidentalis*) spring vs. summer growth provides an example of the type of quantitative data that can be produced from Rapid-Tome sections (Fig. 4M-R). This analysis utilizes the high contrast of autofluorescent cell walls viewed with a GFP filter cube. Image 4M is the raw fluorescent image that is then made binary (4N) for the “analyze particles” command in FIJI to detect and measure the summer and spring xylem vessel area (4Q and 4R, respectfully). The summer growth (4O and 4Q) contains visibly smaller xylem which were determined to be 23% of the total area of the region of interest. The spring growth (Fig. 4P and 4R) however, is comprised of 49% xylem vessels, a more than 2-fold increase over the summer growth.

Figure 5 shows sections taken with the Rapid-Tome generated by students with no prior sectioning experience during an hour-long workshop. Lignin in the sections in Fig. 5 is stained with phloroglucinol-HCL. The ability to make these clear measurements and observations exemplifies the utility of the Rapid-Tome in anatomical investigations in both research, classroom, and demonstration laboratories.

**Figure 5.**
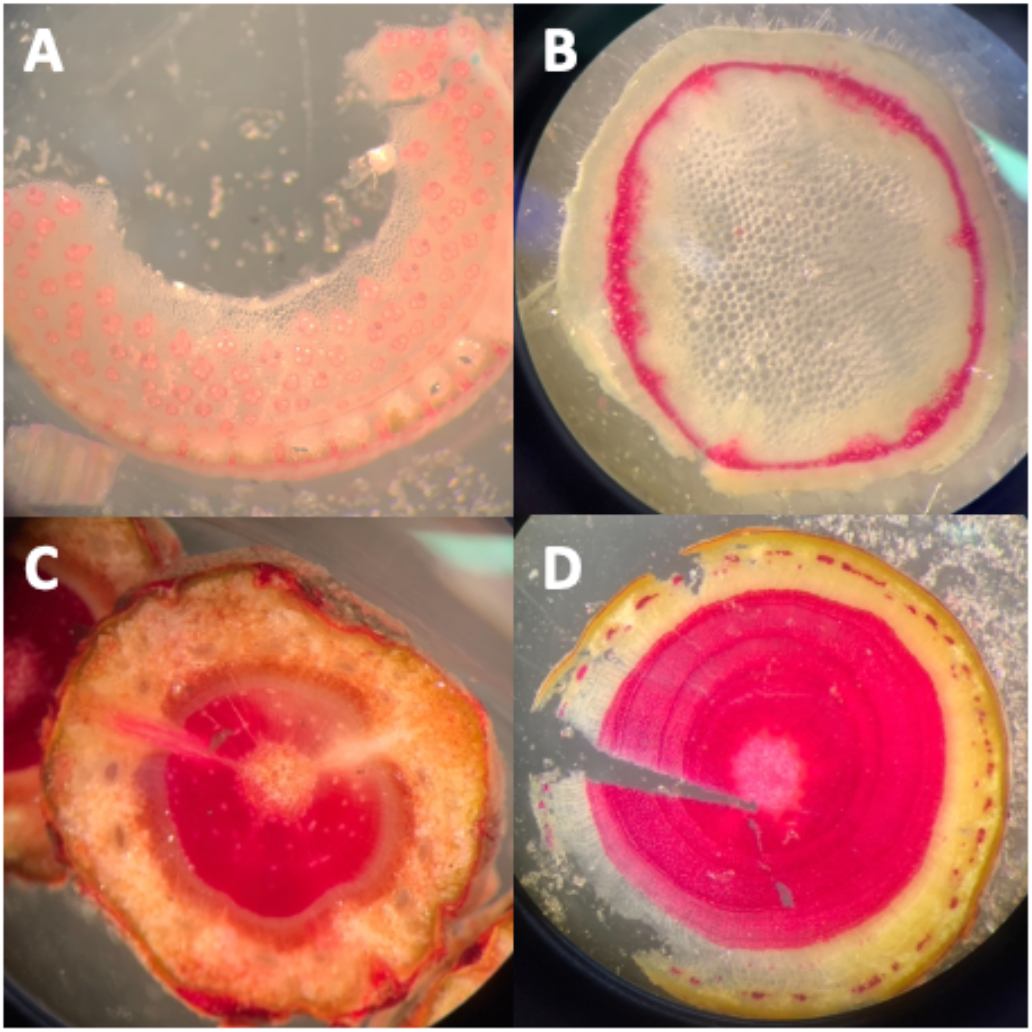
Sections prepared with the Rapid-Tome by students with no prior sectioning experience. The quality of the sections shows that the learning curve with this tool is low. Students were given a protocol and hands-on demonstration of the operation of the Rapid-Tome and had less than an hour of practice to section and stain various samples with phloroglucinol-HCl. (A) Giant reed (*Arundo donax*), (B) Mint *(Mentha piperita L*.) (C) Shore pine *(Pinus contorta)*, and (D) Apple *(Malus domestica)*. Images collected with a cell phone through the eyepiece of a dissection microscope.

Finally, we also developed a method for achieving longitudinal sections that utilizes a carrot to support the sample while sectioning with the Rapid-Tome (Additional File 2). The segment in this example is the soft tissue of an Alocasia petiole which has been pushed through the cut hole of the carrot, which is then trimmed to fit through the opening in the stage of the Rapid-Tome (Additional File 2C and D). Additional File 2 E shows a partially sectioned carrot with stem segment intact and the “grip” that the carrot has to secure the alocasia petiole in place while sectioning. A thin longitudinal section with visible vascular bundle fibers between the dark epidermis (Additional File 2F).

## Conclusions

Microscopic analysis of plant anatomy is a common procedure to study plant structure and function, serves as an integral component of phenotype screening in functional genomics, and is a staple in plant biology coursework. Observations and measurements of plant anatomy rely on high-quality sections for mounting on microscope slides. Section quality depends on the sectioning tool, the researcher, and the qualities of the plant sample. Hand sectioning is adequate for herbaceous samples that lack extensive lignification of cell walls. To section woody and lignified stems, benchtop microtomes can produce extremely thin sections (<100 microns) and have been commonplace in labs for many decades. However, to achieve the quality of section required for analysis, these instruments typically require a lengthy process of resin or wax embedding, are time consuming to set up, possess a steep learning curve, and can be prohibitively expensive. Here we present an updated and improved hand sectioning method that is suitable for hollow-stemmed grasses and similar plant materials. Providing a detailed description of a common method for producing sections of plants for microscopy reduces the amount of time a novice may require achieving high-quality sections of hollow stems.

In addition, we present the design and use of the novel hand sectioning device, the “Rapid-Tome.” This device lowers the learning curve for safely producing high-quality sections from tougher plant materials with highly lignified cell walls such as woody stems and mature, highly fibrous herbaceous plants. Our images and analysis show that the sections produced by the Rapid-Tome are suitable for comparative analyses often conducted with plant anatomy that require high-quality sections for clear anatomical observations. Furthermore, the learning curve of using the Rapid-Tome and the low cost to obtain one lowers the time investment for students and new researchers to produce publication quality sections for imaging and observation even within a classroom setting.

## Methods

### Plant material and collection

For Figure 2, switchgrass internodes were collected from field-grown plants from Columbia, Missouri in September of 2018 before the first killing frost. Each internode segment included 2 cm of internode below the lowest node above soil level and approximately 8 cm of internode above the node. Switchgrass internodes were stored in 50% EtOH and distilled water at 4°C. Sections were made starting 1 cm above the node for no more than 1 cm (2 cm above the node). Prior to sectioning, EtOH-stored samples were rehydrated in 100% distilled water overnight (no less than 12 hours).

Additional switchgrass samples for fresh internode images and root sectioning and imaging were harvested from flowering greenhouse-grown plants under stable, well-watered conditions in August of 2020. Less than 1 cm in diameter Juniper and sycamore twigs < 50 cm from the end of branches were harvested from the University of Oklahoma campus in Norman, Oklahoma in July of 2020. Samples used for the high school student workshop were obtained from the Washington State University greenhouse. These samples were sectioned as fresh samples that were washed gently and stored for less than 24 hours in distilled water.

### Hand Sectioning Method for Hollow-Stemmed Grasses

This portion describes sectioning fresh and or hydrated plant samples. There are several key points to consider when hand sectioning plant materials. Make an initial cut a few mm away from the exact place from which you want to obtain sections, then section your way to the desired location. The initial cut must be perpendicular to the length of the stem and can be made with the stem lying horizontal on a lab bench or cutting mat. Either a blade or hand shears as in Additional File 1E can be used to make the first cut, but a fresh razor blade should be used when sectioning. When sectioning, balance your arms by resting your elbows on table surface or against your body. Keep your hands together and take five or more sections without separating your hands. Every time you separate your hands, you have to reset the subject to achieve thin, uniform sections. Do not use a chopping motion. Slice the stem utilizing the length of the blade. Catch sections with a water droplet resting on top of the blade. Take approximately twice the number of sections you need. Samples will not section well and may bend or collapse while attempting to section them if allowed to partially dry. If partially dried, rehydrate in distilled water or 50% ethanol and distilled water overnight. Below is a more detailed description of this approach with attention to key aspects of the method.

#### Blade considerations

Always use new blades that are rust free and have been stored in a dry place. It makes a big difference which blades are used. We use PTFE coated single edge blades made by GEM, available from Electron Microscopy Supply (item #71970). The silica content in grass stems dulls the cutting edge rather quickly. If the blade does not cut well, use a fresh blade. We typically use a new blade for each new stem or after cutting about 10-20 sections. If blade cost is a concern, it is also possible to reorient the blade in your hands to use a different part of the blade when switching to a new sample or if sectioning is poor. It is important to hold the stem firmly vertical and make a horizontal cutting motion to ensure a flat section is taken. If the stem is not vertical or the blade is not held horizontal the section will be wedge-shaped which interferes with consistent focus in the field of view when imaging. Also, oblique, and wedge-shaped sections can exaggerate the cell lumen area if the blade is not perpendicular to the length of the stem. This is less of an issue with small diameter stems or if cell dimension measurements are not desired.

It is important to slice with the length of the blade rather than a chopping motion to prevent the hollow samples from collapsing. For example, when sectioning a 3 mm diameter stem, approximately 1 cm of the cutting edge of the blade is used, to utilize the blade length. This sectioning approach is like slicing bread, which is easily crushed by a knife, requiring instead a long knife with a long blade to slice rather than chop as you would a rigid carrot. The first section will be too thick and will be discarded. This is also when corrections to the angle of the first cut can be made by making sure that subsequent cuts are perpendicular to the length of the stem.

#### Holding the stem

If the sample is long enough, the length of the stem to be sectioned from can be cut to size to match approximately the length from tip of the researcher’s index finger to the base of the thumb (Fig. 1A). The length will be different depending on the size of the hands. This way the opposite end of the stem can rest steady against the palm while being held by the tips of the thumb and index finger with the index finger slightly above the tip of the thumb (Fig. 1B and 1C).

#### Holding the blade

The blade is held in your dominant hand between the middle finger and thumb with the blade facing toward you and the length of the blade parallel to the blade holding thumb (Fig. 1D). The blade is held firmly and horizontally by the dominant hand with the broadside of the blade pressed firmly downward against the index fingertip holding the stem. The primary movement of the blade comes from the tip of the index finger holding the stem, bending back and forth to advance the blade horizontally to cut the section (Fig. 1G and H).

#### Hand sectioning

When sectioning, the blade moves forward through the stem with the movement of the index fingertip. The dominant hand index finger holding the blade also applies enough pressure to slightly rotate the blade (∼15º) with the pivot point between the dominant hand thumb and middle finger (Fig. 1E and F). The key is to make multiple sections (5-10) without separating the hands from each other. This is to keep the vertical separation of the blade and the cutting surface in as consistent proximity as possible. Holding the blade so that it is resting on top of the cut surface, slightly apply downward pressure (held in balance against opposing pressure of the index fingertip that is holding the stem) and pull the blade backwards so the cutting edge just barely drops off the side of the stem. The downward pressure should be just slightly more than the upward pressure so that the blade height moves down minutely after reaching the edge of the stem’s cut surface. This is to zero in on the level at which the last section was taken and to orient where to cut the next section. This is a key movement to get very thin sections (< 100 μm) and takes practice.

To expose more stem to section without taking your hands apart, advance the stem upwards very slightly by bending backwards the thumb that holds the stem, in between taking sections (Fig. 1I and IJ). The first cut or two get the cutting surface of the stem situated in the right place relative to your hands. The best sections will typically be the 3^rd^ or later once you get the motions dialed in. Move the stem upwards ever so slightly by rocking your thumb slightly backwards after each cut. Plan to cut at least twice the number of sections that are needed.

It is important to keep the sample wet and hydrated during sectioning, touch the end that is being sectioned to water after the initial cut and after ∼10 sections are taken. Dip the cutting edge of the blade into water so a few microliters of liquid are resting on the cutting edge where the sections will be. The water droplet helps to prevent the section from flying off and getting lost and helps lubricate the cut. Once 5-10 sections are taken without separating your hands, collect the sections with forceps or by dipping the blade into water to “wash” them off into a shallow vessel or petri dish.

### The Rapid-Tome 3D-printed hand-held microtome

#### Printing instructions with polylactic acid (PLA) filament

See Additional File 3 for printing directions, Rapid-Tome assembly, and sectioning protocols. Additional Files 4 through Additional File 14 are 3D print files in .stl format. Print all parts in provided orientation for optimal quality. The recommend setting for the filament extruder resolution is “high detail.” We had great success with 0.14 mm layers on a TAZ 6 Lulzbot. On a Snapmaker 2.0 the dimensions were precise with a layer height of 0.24 mm. These are simply the default printer setting for “high detail” and “normal fast print,” respectively. The exact settings required will vary from printer to printer. The default “high quality” setting for any printer should be adequate, though may not be needed if using more expensive or high-quality printers. Higher quality settings and thinner layer heights add print time but will ultimately provide a better tool. Even at high-quality printer settings, all parts should be printable in less than 24 hours. To test the quality attainable at each setting for a printer, start by printing the nut and blade clamp screw. These parts require the highest accuracy and will serve as an accuracy test. The setting at which these parts can accurately be printed and tightly fit together is the maximum print quality needed. Furthermore, these parts are relatively small and can be printed in just a few hours or less. All parts should be printed no less than 15% infill. Add brims to all parts, though a raft would be effective as well if desired but adds print time. Make sure to add supports to all overhangs greater than 70 degrees with at least 20% infill.

#### Assembly instructions

Once printed, remove all supports and brims, etc. so that the pieces resemble those in Fig. 3C and Additional File 1. Though not required, we recommend that you apply spray-on PTFE dry coating lubrication to the contact points on the left and right edges of the stage (Additional File 1G), and within the C-shaped grooves of the sled (Additional File 1O). Non-3D-printed parts required are listed in Table 2 and assembly steps are shown in Additional File 3.

**Table 2.** Non-3D-printed parts for Rapid-Tome.

| Part | # Required | Description |
| --- | --- | --- |
| 1. Bolt (A) | 1 | Machine screw<br>#8-32 x 1.5 inch |
| 2. Nut (A) | 1 | Nut<br>#8-32 |
| 3. Advancement Screw | 1 | Machine screw<br>#8-32 x 4 inch |
| 4. Advancement nut | 2 | Nut<br>#8-32 |
| 5. Drawer pull knob | 1 | Large diameter and flat, must fit the #8-32 machine screw |
| 6. Washer | 1 | Stainless steel ½ inch inner diameter, 1 3/8 inch outer diameter flat washer*. |
| 7. Self-adhesive High density weather strip | 1 x .25 inch strip | Cut to size from: Duck Brand Foam Weatherstrip Seal for Small Gaps (found at Walmart), must be dense foam. |
| 8. Razor blades | Box of 100 | Single Edge, GEM “PTFE” Coated<br>#71970<br>https://www.emsdiasum.com/microscopy/products/preparation/blades.aspx#71970 |
| **Additional Materials** |  |  |
| 9. Blaster Advanced Dry Lube with Teflon | 1 can | Widely available.<br>Model # 16-TDL |
| 10. LOCTITE® Super Glue Control Gel 4 gr | 1 | Widely available.<br>234790 en-US |
| 11. Screw driver | 1 | Match machine screws |

\* Washers are available here: https://www.amazon.com/Stainless-Steel-Flat-Washers-USS/dp/B094D3X5RT?th=1.
Note: Zinc and galvanized washers of this size are available in store at Home Depot, not Lowe's. Of the two, zinc is better because it is without the galvanized (usually zinc) coating that is scratched off by the blade, though rather roughly and can cause sectioning issues and bumps in the washer surface. The zinc will work for a short time but is softer than steel and so will accumulate nicks from the steel razor blade, leading to damage of subsequent razor blades and poor sectioning. Therefore, stainless steel is the best option.

First, install the advancement screw by placing one of the nuts over the screw about 5 mm and sliding the screw into the opening at the base of the handle (Additional File 1A). With a screwdriver, turn the screw until the end reaches the other side of the hexagonal seat on the bottom of the handle (Additional File 1B). Place a nut into the base of the handle and turn the screw into it. If the nut is not flush with the base of the handle (Additional File 1C), back the screw out of it, lift the nut out carefully and rotate 180 degrees and put back into the seat for the nut. This rotates the bite point for the screw within the nut so that the nut should be held flush with the bottom of the handle (Additional File 1D). The two nuts should both be firm against and sandwiching the base of the handle, without any play. This is crucial to insure proper control on the slider and sample advancement during sectioning.

After advancing the screw 1 cm through the bottom of the handle, attach the drawer knob and firmly tighten the screw all the way into the knob. Place a slider into the slider groove (Additional File 1E). Multiple sliders can be stacked according to the length of the sample segment or if a shorter advancement screw is used. Two sliders can be stacked for short samples (Additional File 1F). Line up the pegs on the top of the handle with the holes on the underside of the stage, thread the screw into the opening through the stage, place nut over the screw on the back of the handle and tighten down firmly (Additional File 1G).

Install the ½ inch washer by gluing it into place with epoxy. Note that the opening of the sled and the washer edge must align, with neither overhanging the other (Additional File 1I). Apply a 3mm wide bead of LOCTITE® Super Glue Control Gel around the stage opening to hold it in place. Apply the weather stripping to the blade clamp (Additional File 1K). Weather stripping may need to be replaced occasionally.

Insert the printed nut into the slot on the back of the sled, (Additional File 1L). Place the printed washer over the screw and thread it through the blade clamp. Place a razor blade into the sled ensuring the notches in the blade fit around the tabs in the sides of the blade bed (Additional File 1L). Attach the blade clamp and tighten down as far as possible with the printed screw, approximately 2.5 turns (Additional File 1M). Install the blade guard by placing the tip of the guard into the slot beneath the Rapid-Tome label, then press the squared column into the opening on the top of the sled (Additional File 1N, H, and J).

When ready to section, remove the blade guard by pulling the squared grip indention. Place the blade guard into the holster, slight bending is required to ensure it stays in place. Insert the tip first, then press the squared column downward into place. Slide the sled onto the stage nearly halfway. The blade may need to be gently pushed from beneath to clear the front edge of the washer the first time the blade is used.

#### Cross sectioning of cylindrical samples

As described above, cut the specimen to approximately hand-length before using the Rapid-Tome. Place the specimen into the Rapid-Tome handle groove and firmly hold with your non-dominant-hand’s thumb against the back of the handle. Turn the advancement knob with your dominant hand to make the top of the segment just above the surface of the washer. Note that the sample must be held firmly against the back of the handle groove. Woody twigs that are not completely straight will need to be rotated until firmly against the back wall, as required for a clean cut. If this is not followed, the section may not be fully freed from the specimen and the specimen may split. For brittle or fragile samples, we found that two layers of stretched parafilm help to prevent splitting or crushing. Once the specimen is cleanly positioned, with the dominant hand, place your thumb on the front of the sled and your forefinger on the back of the stage.

Make the first cut by sliding the sled forward by pressing your thumb and forefinger together. Remove section with tweezers and immediately place into water. If using the optional section catcher, flick sections into the catcher with a small paintbrush or forceps. The catcher should contain some drops of water to prevent sections from drying out.

Once a section is made, the forefinger of the non-dominant hand can push the sled back to the start position. Turn the advancement knob 180 degrees. The amount rotated can be adjusted according to the sample and the desired section thickness. Advance the sled again toward your sample. Avoid moving the sample in any way in between sections, other than by the advancement screw. Hold the sample in place and take five to ten sections at a time without removing your hands or resituating the sample (other than vertical advancement). This helps get the best sections possible. If the blade is dulled or bends and catches on the washer as can happen with tough samples, try to keep hold of the sample in the handle with your left hand, using only your right hand to unscrew the blade clamp and replace with a new blade.

#### Longitudinal sectioning of cylindrical samples

The method to achieve longitudinal sections with a carrot to quickly support the tissue is an adaptation of a tried-and-true approach that utilizes the structural integrity and cut-ability of the widely available root vegetable (Additional File 2). For this approach, a hole is first cut through the carrot perpendicular to its long axis with a drill bit that matches the diameter of the plant tissue or is slightly smaller (Additional File 2A and 2B).

### Staining

Acriflavine hydrochloride was used in Figure 4 G-L at 0.1% w/v as described in [21]. Crystal violet was used on the Juniper sections at 0.1% w/v [22]. Toluidine blue O was used for the switchgrass roots in Fig. 4D-F at 0.02% w/v in distilled water as described in [23]. Phloroglucinol-HCL is widely used to visualize lignin in cell walls indicated by a bright red color used in Fig. 5 [24].

### Image collection

Unless otherwise stated, all brightfield illumination microscopy completed on a Nikon Eclipse Ni microscope and a Nikon Ds-Qi1Mc camera with an LED light source. Scale bar applied in FIJI [25]. Fluorescent image in Figure 4 M captured with the same microscope and a Nikon DS-Qi2 monochrome camera.

## Supporting information

Rapid-Tome Supplemental Files

## Additional Files List

1. Additional file 1. Additional Figure 1. Assembly of the Rapid-Tome.
2. Additional file 2. Additional Figure 2. Longitudinal sections can be made utilizing a hole cut through a carrot.
3. Additional File 3. Quick protocols for Rapid-Tome printing, assembly, and sectioning.
4. Additional File 4. All Rapid-Tome Complete File Set.stl
5. Additional File 5. Rapid-Tome Blade Guard.stl
6. Additional File 6. Rapid-Tome Clamp.stl
7. Additional File 7. Rapid-Tome Handle.stl
8. Additional File 8. Rapid-Tome Nut.stl
9. Additional File 9. Rapid-Tome Screw.stl
10. Additional File 10. Rapid-Tome Section Catcher.stl
11. Additional File 11. Rapid-Tome Sled.stl
12. Additional File 12. Rapid-Tome Slider.stl
13. Additional File 13. Rapid-Tome Stage.stl
14. Additional File 14. Rapid-Tome Washer.stl

## Ethics approval and consent to participate

Not applicable

## Consent for publication

Not applicable

## Availability of data and materials

Files in .stl format for the 3D-print parts are available as Additional Files and will be made available through the plant phenomics website (3Dplantphenomics.org) and file hosting through a file repository on GitHub. Print files will also be made available on the leading online 3D printing design repository, Thingiverse.

## Authors Contributions

DT developed the hand-sectioning method and designed, optimized printing, and tested the Rapid-Tome. DT and LEB wrote the manuscript. JR conducted additional testing of the Rapid-Tome and made comments on the manuscript.

## Acknowledgements

Bobby Reed at the Innovation @ the EDGE at the OU Bizzell Memorial Library made this project possible by printing the numerous iterations for testing leading to the final design. Jordan Rainbow conducted additional printing and sectioning testing of the Rapid-Tome. Dan Manwaring at the WSU Fine Arts Center also provided printing services and insight into printing optimization. Andrei Smertenko and Taras Nazarov led the workshop with high school students that produced Figure 5 with support from NSF-CAREER award #1751204.

## Author’s information

ORCID IDS: David J. Thomas 0000-00001-9443-0527, Laura Bartley: 0000-0001-8610-7551

