## Supplementary material for "The Rapid-Tome, a 3D-Printed Microtome, and an Updated Hand-Sectioning Method for High-Quality Plant Sectioning": Rapid-Tome Supplemental Files

#### Additional Files and Figure Legends:

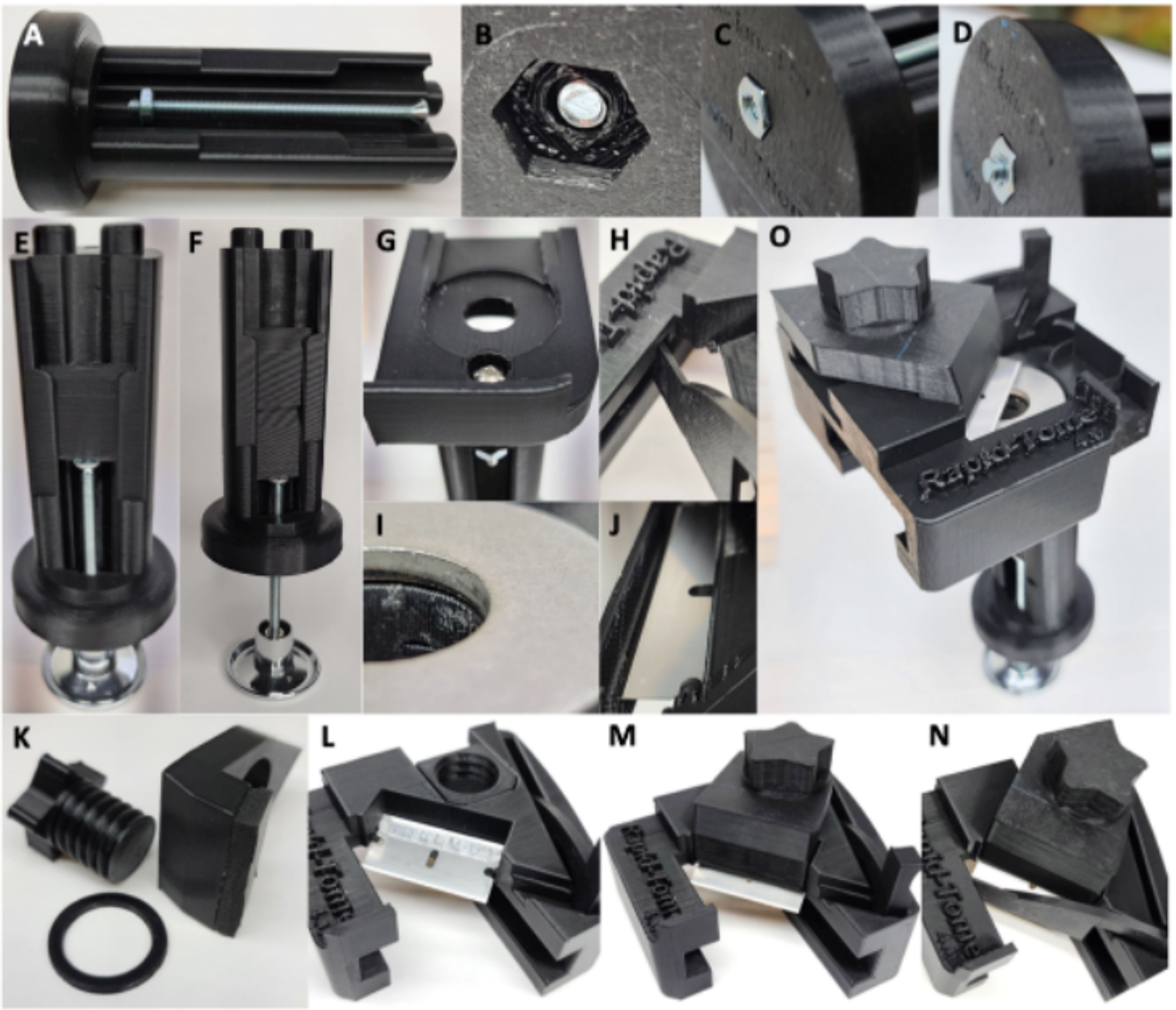


**Additional File 1. Assembly of the Rapid-Tome.** (A-D) The advancement bolt is installed first. (E-G) The slider is placed within the grooves of the handle before the stage is attached. (F) Two sliders can be stacked if a short sample is to be sectioned. (H-J) The washer and blade guard are attached. The blade clamp (K) is crucial and holds the blade in place (L-N). (O) Fully assembled Rapid-Tome. Sliding the sled into place over the stage against the inclined washer flexes the blade and ensures a flat cut is made. All plastic parts are 3-D printed with PLA.

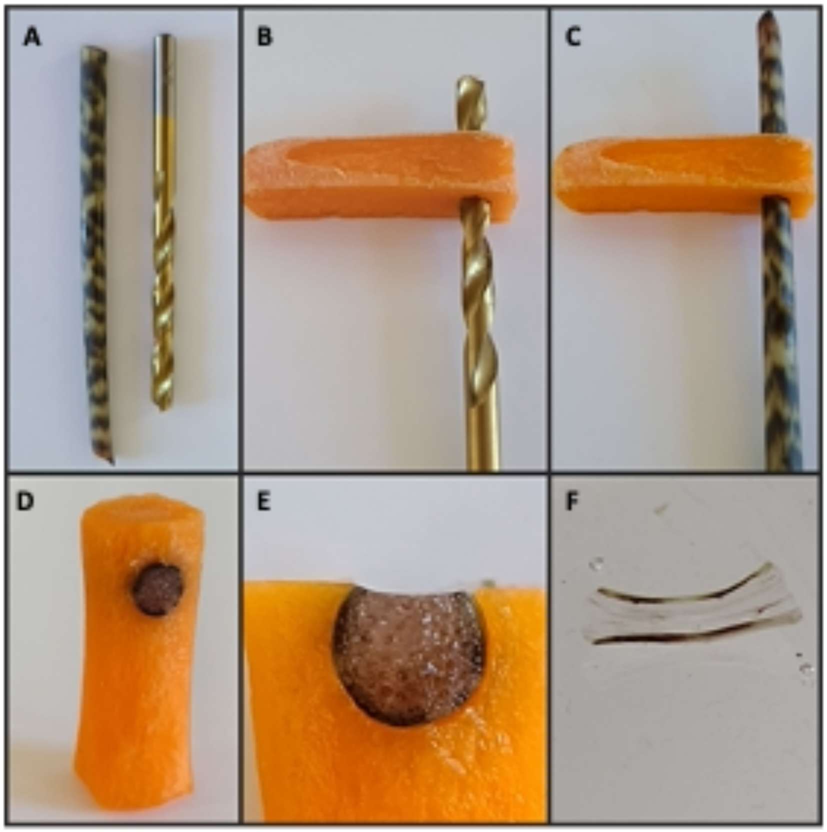


**Additional File 2 . Longitudinal sections can be made utilizing a hole cut through a carrot.** (A) Select a drill bit that is the same diameter or slightly smaller than the *Alocasia* sp. stem segment. (B) Drill a hole through a carrot gently by hand. (C) Push the stem segment into the carrot until snug. (D) Use a razor blade to remove excess stem segment and carrot so that the carrot fits into the Rapid-Tome stage hole. (E) The carrot effectively holds the stem in place during sectioning. (F) Thin sections are possible even with fragile stem tissue.

**Additional File 3. Directions for printing and assembling the Rapid-Tome**

**Protocols:**

### Rapid-Tome

#### 3D printed Rapid-tome parts:

Only one copy of each component below is required except for the slider if short samples are to be sectioned in which case, two sliders are recommended.

- Handle
- Stage
- Sled
- Blade clamp
- Blade clamp Screw
- Blade clamp washer
- Blade clamp nut
- Blade guard
- Slider
- Section catcher

#### Printing instructions

1. Print all parts in provided orientation for optimal quality.
2. We recommend setting the filament extruder resolution to be high detail. We had great success with 0.14 mm layers on a TAZ 6 Lulzbot. On a Snapmaker 2.0 the dimensions were precise with a layer height of .24 mm. These are simply the default printer setting for "high detail" and "normal fast print", respectively. The exact settings required will vary from printer to printer. The default "high quality" setting for any printer should be

adequate, though may not be needed. The filament we test printed with exclusively is PLA, polylactic acid, though ABS, Acrylonitrile Butadiene Styrene, and others would work as well with slightly modified printing specifications according to filament manufacturer's recommendations.

3. To test the quality capable at each setting for a printer, start by printing the nut and blade clamp screw. These parts require the highest accuracy and will serve as an accuracy test. The settings at which these parts can accurately print and tightly fit together is the max print quality needed for all parts. Furthermore, these parts are relatively small and can be printed in just a few hours or less.
4. All parts should be printed no less than 15% infill.
5. Add brims to all parts, though a raft would be effective as well if desired but adds print time.
6. Make sure to add supports to all overhangs over 70 degrees with at least 20% infill, or as recommended by your printer.
7. These models would likely work in resin printers as well, though we did not test that or have any printing recommendations for that type of printer.
8. NOTE: trying to print at lower quality to save time will result in frustration and in fact waste your time. A low-quality print will make the contact points incorrect and render the tool unfunctional. If the pieces do not fit together as described, adjust your printer settings to reduce layer height and or slow down the print speed.

**<u>Assembly instructions</u>**

1. Remove all supports and brims, etc.

2. Install advancement screw: place one of the nuts over the screw about 5 mm as seen in Figure S3 A and slide into the opening at the base of the handle. With a screwdriver, turn the screw until the end reaches the other side of the hexagonal seat on the bottom of the handle. Place a nut into the base of the handle and turn the screw into it. If the nut is not flush with the base of the handle, as seen in Figure S3 C, back the screw out of it, lift the nut out carefully and rotate 180 degrees to adjust the placement of the threads within the nut and put back into the seat for the nut. This rotates the bite point for the screw within the nut so that the nut should be held flush with the bottom of the handle as seen in Figure S3 D. The screw should not have any play or rattle around. In other words, the two nuts should both be firm against and sandwiching the base of the handle. This is crucial to insure proper control on slider and sample advancement during sectioning.
3. After advancing the screw ~1 cm through the bottom of the handle, attach the drawer knob and firmly tighten the screw all the way into the knob.
4. Place a slider into the slider groove. Note: the number of sliders can be adjusted according to the length of the sample segment or if a shorter advancement screw is used.
5. Line up the pegs on the top of the handle with the holes on the underside of the stage, thread the screw into the opening through the stage, hole A, and place nut over the screw on the back of the handle and tighten down firmly.
6. Install the ½ inch washer by gluing it into place with epoxy. Note that the opening of the sled and the washer edge must align, with neither overhanging the other (Additional File 1I). Apply a 3mm wide bead of LOCTITE® Super Glue Control Gel around the stage opening to hold it in place.
7. Insert the printed nut into the slot on the back of the sled seen in Figure 3 B.

8. Place the printed washer over the screw and thread it through the blade clamp.
9. Place a razor blade into the sled ensuring the notches in the blade fit around the tabs in the sides of the blade bed Figure S3 L. Attach the blade clamp and tighten down as far as possible with the printed screw, (approx. 2.5 turns), Figure S3 M.
10. Install the blade guard by placing the tip of the guard into the slot beneath the Rapid-Tome label, then press the squared column into the opening on the top of the sled Figure S3 N.
11. When ready to section, remove the blade guard by pulling the squared grip indention. Place the blade guard into the holster, slight bending is required to ensure it stays in place. Insert the tip first, then press the squared column downward into place.
12. Slide the sled onto the stage nearly halfway but not to where the blade covers the opening in the stage Figure S3 O. The blade may need to be gently pushed from beneath to clear the front edge of the washer the first time the blade is used.

**<u>Sectioning</u>**

**Cross sectioning of cylindrical samples:**

1. Cut the sample to length before using the Rapid-Tome. This can be done with shears or a razor blade on bench top,
2. For right-handed people:
3. Place the segment into the groove and firmly hold with your left-hand thumb against the back of the handle. Turn the advancement knob with your right hand to make the top of the segment just above the surface of the washer. Note: the sample must be firmly against the back of the handle groove. Woody twigs that are not completely straight will need to be rotated until firmly against the back wall that is required for a clean cut. If this is not followed, the section will not fully cut from the sample segment and the sample will likely

split. For grasses and herbaceous samples, 2 wraps of stretched parafilm help to prevent splitting as well as crushing.

4. With right hand, place your thumb on the front of the sled as seen in Figure 1 and your forefinger on the back of the stage.
5. The left-hand forefinger is used to push the sled back to the start position.
6. Make the first cut by sliding the sled forward by pressing your thumb and forefinger together.
7. Remove section with tweezers and immediately place into water
8. If using the section catcher, flick sections into the catcher with a small paintbrush or forceps. Catcher should contain some drops of water to prevent samples from drying out.
9. Turn the advancement knob 180 degrees. The amount rotated can be adjusted according to your sample and desired section thickness.
10. Advance the sled again toward your sample. Note: do not move the sample in any way in between sections, other than by the advancement screw.
11. If the blade is dulled or bends and catches on the washer as can happen with tough samples, keep hold of the sample in the handle with your left hand, using only your right hand to unscrew the blade clamp and replace with a new blade.

**Important sectioning tips**:

1. Hold the sample in place and take 5 – 10 sections at a time without removing your hands or resituating the sample (other than vertical advancement). This is to help get the best sections possible.
2. Apply spray on PTFE dry coating lubrication to the contact points on the stage and the sled.
